# Expanding the human T cell receptor germline reference through population-scale pangenome analysis using the gAIRR Suite version 2

**DOI:** 10.64898/2026.11.14.766613

**Authors:** Yu-Hsuan Yang, Yu-Hui Lin, Kuan-Ta Huang, Mao-Jan Lin, Jacob Shujui Hsu, Pei-Lung Chen

## Abstract

The germline gene set encoding the adaptive immune receptor repertoire (gAIRR) defines the allele space from which the expressed repertoire (exprAIRR) is assembled, so an incomplete gAIRR reference propagates errors into T cell receptor (TR) annotation and genotyping and is also a barrier to studying inherited TR variation in its own right. Human TR gAIRR diversity remains incompletely characterized because the loci resist short-read resolution and because locus-resolved cohorts have been small. We developed gAIRR Suite version 2, in which each gAIRR-annotate query carries 50 bp of germline sequence on either side of the allele core and a call is reported only when the assembly supports the core together with both flanks, and in which gAIRR-call genotypes from Element AVITI libraries whose 500-bp Read 1 spans a complete V core with its flanking anchors. Applying this framework to Human Pangenome Reference Consortium Release 2 (464 haplotype-resolved assemblies from 232 individuals across five superpopulations), we identified 531 novel TR alleles (486 TRV, 45 TRJ), each observed in the targeted-sequencing reads of at least one carrier, of which 298 were absent from our 47-individual Release 1 catalogue. In 216 individuals, targeted genotyping with gAIRR-seq and gAIRR-call recovered 99.985% of the 61,628 evaluated alleles found in the corresponding assemblies, with a precision of 99.995%. Release 2 added few common alleles but many rare ones: 407 of the 531 were restricted to a single superpopulation, and African-ancestry individuals accounted for 51.1% of these (46.0% ± 2.1% after downsampling to equal group size) while comprising 30.2% of the cohort. Population-scale pangenome analysis therefore extends the TR gAIRR reference chiefly by recovering variation that smaller or less diverse cohorts would miss.

## Introduction

The adaptive immune receptor repertoire (AIRR), comprising the T cell receptors (TR) and immunoglobulins (IG), determines the range of antigens an individual can recognize [1,2]. These receptors are generated during lymphocyte development by V(D)J recombination [3,4], with further diversity introduced at the junctions through nucleotide addition and deletion [3,5]. Two entities are therefore involved, which we distinguish as previously [6]: the germline gene set encoding AIRR (gAIRR), inherited and shared by every cell of an individual, and the expressed AIRR (exprAIRR), the somatically rearranged repertoire to which the term "AIRR" ordinarily refers. Much of exprAIRR nonetheless remains germline encoded: V segments encode the framework regions and CDR1/CDR2, whereas CDR3 is formed at the junction [7]. exprAIRR is therefore shaped by the gAIRR allele set from which it is derived [6,7].

Accurate exprAIRR profiling consequently depends on the completeness of the gAIRR reference used for V(D)J annotation [2,8]. This is particularly important for TR because, unlike IG genes in antigen-experienced B cells, rearranged TR genes do not undergo somatic hypermutation: outside the junction, a TR sequence is expected to match its inherited allele, and mismatches reflect either uncatalogued germline variation or technical error [6,8]. When an allele is absent from the reference, reads are misassigned to related alleles, producing incorrect gene and allele calls that bias downstream measurements such as gene usage [8,9]. These errors are especially relevant for populations underrepresented in current gAIRR reference resources [10].

gAIRR variation is also a heritable determinant of immune phenotypes in its own right, independently of its role as an annotation reference. Non-coding variation in recombination signal sequences (RSS) can modulate recombination efficiency [11–13], and coding TR polymorphisms can alter antigen recognition and have been associated with clinically relevant immune phenotypes [7,14,15]. A recent population-scale analysis further showed that inherited TR variation carries signatures of natural selection and influences thymic fate commitment and cell-surface receptor abundance [7]. A more complete TR gAIRR catalogue is therefore needed for both purposes.

Human TR germline diversity nonetheless remains incompletely characterized for two reasons. First, the loci are difficult: the interdigitated TRA/TRD locus on chromosome 14 and the TRB and TRG loci on chromosome 7 contain extensive segmental duplications and near-identical paralogues that short-read sequencing cannot resolve [7,16,17]. Locus-resolved genomic studies were until recently limited to cohorts of tens of individuals [6,7,17], restricting discovery of rare and population-enriched alleles [6,10]. Second, existing reference resources remain incomplete in sequence context. IMGT/GENE-DB provides the principal curated reference for human TR germline genes and alleles [18], but some catalogued alleles lack complete 5′ and/or 3′ flanking sequence [6]. These gaps limit characterization of RSS and complicate precise allele-boundary definition. Community resources such as VDJbase and OGRDB broaden access to inferred alleles [19], but flanking context requires high-quality genomic sequence [6,17].

Long-read, phased whole-genome assemblies now provide it: the Human Pangenome Reference Consortium (HPRC) offers ancestry-diverse, haplotype-resolved assemblies with the contiguity and accuracy needed to resolve complex loci [20,21]. Several tools annotate germline TR segments from assemblies. gAIRR-annotate aligns IMGT-defined core V-, D-, and J-REGION sequences to an assembly and reports those differing from the reference as novel candidates [22]; as in IMGT nomenclature [18], allele identity is defined by the core alone. Flanking sequence is not part of the allele definition, but it delimits segment boundaries and is the substrate for RSS analyses. In version 1, gAIRR-annotate detected alleles from the core alone and recovered flanking sequence afterwards; whether a core-identical call represented a full-length germline allele was resolved by a separate comparison step, which required manual inspection and a hand-curated set of substitute flanks for IMGT entries lacking them [6,22]. The gAIRR Suite pairs gAIRR-annotate with an independent experimental route, gAIRR-seq targeted capture and gAIRR-call genotyping [22]. Applied to 47 HPRC Release 1 individuals, it identified 335 novel TR alleles, increases of 91.6% (TRV) and 30.9% (TRJ) over IMGT [6]. That analysis established the approach, but two separate limits stood in the way of scaling it. Rare and population-enriched alleles are largely absent from small cohorts, so recovering them requires more individuals rather than a more sensitive method — a limit of the 47-genome scope, not of the pipeline. The manual flank comparison was the opposite: tractable at 94 haplotypes, impractical at 464. Release 2 therefore required both a larger cohort and a version of the suite in which flanking context is applied automatically to every query.

We therefore developed gAIRR Suite version 2. Each gAIRR-annotate query carries, for every gene, 50 bp of germline sequence on either side of the core, and a call is reported only when the assembly supports the core together with both flanks. Flanking sequence thus anchors the core — particularly for the short D and J segments — without entering the allele definition, yielding more conservative calls. On the experimental side, gAIRR-seq version 2 libraries sequenced with a 500-bp Read 1 allow single reads to span a complete V core with its flanking anchors, so gAIRR-call version 2 genotypes from full-span read evidence rather than phasing variants across partial reads.

Here we apply this framework to HPRC Release 2, comprising 464 haplotype-resolved assemblies from 232 individuals spanning five superpopulations [21]. We identify 531 novel TR alleles (486 TRV and 45 TRJ), of which 298 (277 TRV and 21 TRJ) were not reported in our Release 1 analysis. We assess this expanded catalogue through cross-release reproducibility, recurrence across individuals and gAIRR-seq/gAIRR-call genotyping, and we characterize allele frequency, population distribution and flanking sequence context. Together, these results strengthen the foundation for gAIRR genotyping, exprAIRR profiling, and studies of the immunological consequences of inherited TR variation.

## Methods

### Allele definitions and reference database

As in our Release 1 study [6], an allele was defined by the nucleotide sequence of the IMGT-defined V-REGION, D-REGION, or J-REGION, with boundaries assigned according to IMGT conventions [18]. Flanking sequences were defined as the unarranged germline genomic sequence immediately upstream and downstream of these regions. For each allele call, 50- and 200-bp flanking sequences were extracted on both sides. RSS were treated as part of the flanking genomic context, so variation within an RSS did not contribute to allele identity.

Novelty was assessed against IMGT/GENE-DB [23] (program version 3.1.43; data accessed on 20 March 2026). The reference set comprised 519 human TR alleles (412 V, 6 D, and 101 J) across all functionality classes. Because allele lookup in gAIRR-annotate is gene-specific and some IMGT alleles are identical across genes, candidate novel alleles were additionally compared by exact sequence with all 519 reference alleles, regardless of gene.

### HPRC Release 2 cohort and genomic DNA

We analyzed 464 haplotype-resolved assemblies from 232 individuals in HPRC Release 2 [21]. Release 2 included 46 of the 47 individuals analyzed in our Release 1 study, excluding HG03492. Superpopulation assignments followed the 1000 Genomes Project panel [24] and comprised 70 African (AFR), 51 East Asian (EAS), 44 Admixed American (AMR), 36 South Asian (SAS), and 31 European (EUR) individuals, with HG002 assigned to EUR and HG005 to EAS as in Release 1 [6], and NA21309 (Maasai) and HG06807 (African American), whose populations are not part of the 1000 Genomes panel, assigned to AFR.

Genomic DNA for targeted sequencing was obtained from the NHGRI Sample Repository at the Coriell Institute (https://www.coriell.org) and was extracted from Epstein–Barr virus-transformed B lymphoblastoid cell lines.

### Assembly-based allele discovery with gAIRR Suite version 2

TR alleles were identified independently in each of the 464 haplotype assemblies using gAIRR-annotate from gAIRR Suite version 2, a rewrite of the Release 1 workflow [6,22] in which the previous BWA- and SPAdes-based steps were removed. Whereas version 1 detected alleles from the core alone and recovered flanking sequence afterwards, version 2 first uses germline flanking sequence to locate the core of each gene in the assembly and then identifies the allele by comparing that core with the alleles of the same gene (Figure 1a).

**Figure 1.**
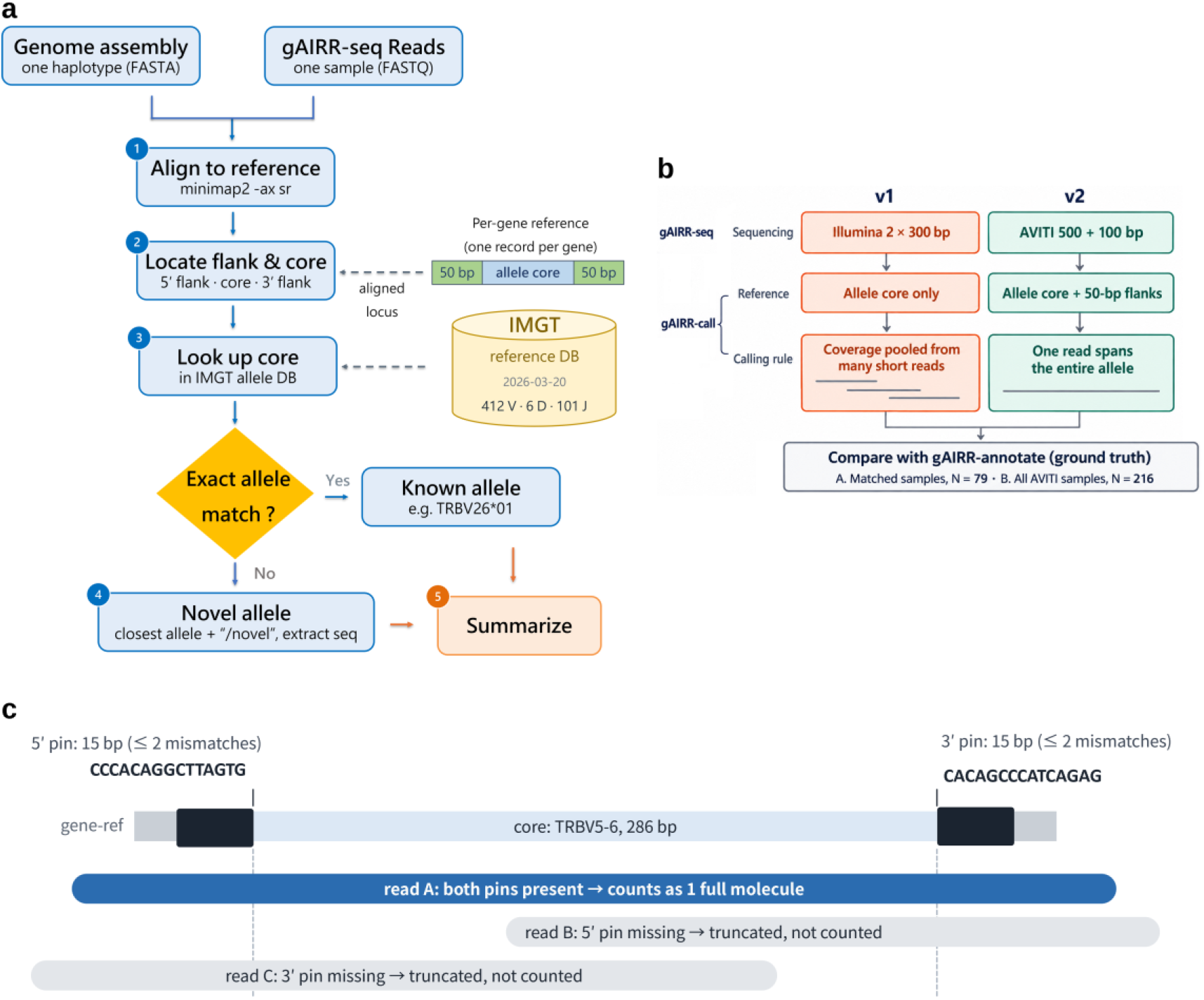
gAIRR Suite version 2 and the benchmarking design. (a) Workflow shared by gAIRR-annotate and gAIRR-call. A haplotype assembly (FASTA) or the gAIRR-seq reads of one sample (FASTQ) are aligned with minimap2 (-ax sr) to per-gene references, each consisting of an allele core with 50 bp of germline flank on either side (1); the flanks locate the core in the aligned locus (2), and the core is looked up in the IMGT/GENE-DB allele set used in this study (accessed 20 March 2026; 412 V, 6 D and 101 J alleles) (3). An exact match is reported as a known allele; otherwise the core is reported as a novel allele, labeled with its closest allele and the suffix /novel, and its sequence is extracted (4). Results are then summarized for each haplotype or sample (5). (b) gAIRR-seq and gAIRR-call version 1 and version 2. Version 1 sequenced Illumina 2 × 300-bp reads, used allele cores alone as the reference and pooled coverage across many short reads; version 2 sequences AVITI reads (500-bp Read 1, 100-bp Read 2), uses allele cores with 50-bp flanks and calls an allele from single reads that span it entirely. Both were compared with gAIRR-annotate genotypes from the same individuals’ assemblies, in the 79 individuals sequenced with both versions (A) and in all 216 individuals sequenced with version 2 (B). (c) Read admission in gAIRR-call version 2, shown for TRBV5-6 (286-bp core). A read counts as one full molecule only if it carries both 15-bp inner flanks (pins; at most two mismatches each), and the core is then taken from the read’s own sequence between the pins; reads missing either pin are truncated and not counted.

For each gene, a single reference was built from the core of a canonical allele (*01 for most genes) with 50 bp of germline flank on each side and aligned to the assembly with minimap2 [25] (version 2.31-r1302) using -ax sr --end-bonus 50 -N 100 --secondary=yes. Seeding parameters were set per locus from the median reference length: -k 15 -w 7 for V-like references (≥250 bp), -k 13 -w 5 for J-like references (130–250 bp), and -k 13 -w 5 -n 1 -m 10 for D-like references (<130 bp), the last allowing a single short seed to chain into a valid alignment. Along each alignment, the assembly bases aligned to the two flank segments were taken as the flanks of the locus and the bases between them as its core, so that the core position was set by the germline flanks while the core itself could differ in length and sequence from the canonical allele. D segments were extracted at the IMGT-defined D-REGION boundaries, without the adjacent RSS heptamers included in version 1.

The extracted flanks were then compared with the germline flanks of the reference, with divergence defined as edit distance divided by flank length, and hits diverging by more than 0.20 on either side were discarded before allele assignment. This removes V(D)J-rearranged loci, in which the RSS-containing flank is replaced by junctional sequence, while tolerating natural germline variation: in the Release 1 flank catalogue [6], germline flanks diverged from the reference by at most 0.16. The filter was applied only to alignments without terminal clipping; a partially aligned hit with a divergent flank was discarded only if another gene whose flank passed the threshold covered the same locus.

Where alignments from several genes overlapped the same locus, the locus was first assigned by flank: a gene was set aside if its flank divergence exceeded that of the best-matching gene by more than 0.10, so that paralogues with near-identical cores were distinguished by flank. Each core was then compared with the alleles of its gene and assigned an allele when it matched that allele exactly. A core matching none of the alleles of its gene was classified as a novel candidate and assigned to its closest allele. Cores sharing less than 90% sequence identity with their closest allele were left unassigned. Where more than one gene remained at a locus after the flank comparison, the gene whose allele matched the core most closely was retained.

Three pairs of reference alleles have identical core sequences: TRBV24-1*02 and TRBV24/OR9-2*01, TRGJ1*02 and TRGJ2*01, and TRBV6-2*01 and TRBV6-3*01. The first two pairs were distinguished by their gene-specific germline flanks. TRBV6-2 and TRBV6-3 are identical across both 50-bp flanks as well; their hits were therefore reported jointly and then assigned from the adjacent annotated gene, based on the conserved gene order TRBV4-2–TRBV6-2–TRBV3-2–TRBV4-3–TRBV6-3–TRBV7-2.

### Novel allele naming and integration into gAIRR Suite

Novel alleles were named as in our Release 1 study [6]. The novel allele sequences from all 464 haplotypes, written by gAIRR-annotate to one FASTA file per haplotype, were pooled, and IgLabel [26] assigned identical sequences to the same provisional four-character identifier. The identifier replaced the /novel suffix that gAIRR-annotate attaches to the closest reference allele (for example, TRBV10-2*01/novel became TRBV10-2*01_V32K).

As in Release 1, the named alleles were then added to the gAIRR Suite allele database. Unlike version 1, which had a single database, version 2 keeps two: material_original, containing only the IMGT/GENE-DB reference set, and material_updated, which adds the novel alleles named here. The database is selected for each run and recorded in its log. Allele discovery and genotyping in this study used material_original, so that novelty was assessed against IMGT/GENE-DB alone. gAIRR-annotate was then rerun on the same assemblies against material_updated to confirm that every novel allele was recognized under its assigned name. Keeping both databases allows new data to be analyzed against either one and the results to be compared directly. All TRV and TRJ alleles observed in the 464 haplotypes, including the 531 novel alleles retained in the catalogue (see Results), are listed with the haplotypes carrying them in Supplementary Table S1.

### Flanking sequence database

For every allele copy annotated by gAIRR-annotate on the 464 haplotype assemblies, the allele sequence was extracted together with 50 bp or 200 bp of flanking sequence on each side, in the orientation of the allele; for D genes the flanks were measured from the D-REGION. Copies of the same allele with identical flanked sequences formed one group, listing the haplotypes that carry it, and the groups of each allele were numbered by the number of haplotypes. The database covers the IMGT alleles and the 531 novel alleles of the catalogue. Copies whose flank was incomplete in the assembly were left out of the affected table, as were flank contexts that contained a somatic V(D)J recombination joint of the cell-line DNA instead of germline sequence. A context was considered to carry such a joint when it was carried by at most two haplotypes, one of its flanks differed from that of every more common context of the same gene by more than 10% of the flank length, and the divergent part contained a segment of a gene that can recombine with it (at least 8 bp of a D-REGION, or at least 20 bp of another V, D or J gene) that was absent from the predominant context of the allele (Supplementary Table S2).

### Targeted sequencing (gAIRR-seq version 2) and genotyping (gAIRR-call version 2)

gAIRR-seq enriches fragmented, adapter-ligated genomic DNA with approximately 120-bp biotinylated probes designed primarily against IMGT-defined V-REGION, D-REGION, and J-REGION sequences [6,22]. For D and J segments shorter than the probe, the probe extended into adjacent germline sequence and could include the RSS.

In gAIRR-seq version 1, libraries were sequenced with Illumina 2 × 300-bp paired-end reads, a configuration that frequently failed to yield individual reads spanning a complete V core [6]. In version 2, libraries were sequenced on the Element AVITI platform in an asymmetric 600-cycle configuration with a 500-bp Read 1 and a 100-bp Read 2 (Figure 1b). Because gAIRR-call version 2 counts a read only when it carries a complete allele core together with 15 bp of germline flank on each side, the required read length is set by the TRV genes, whose cores are approximately 280 bp: a 300-bp read can meet this requirement for only 10 of the 155 TRV gene references (6%), whereas a 500-bp read meets it for all of them, and every TRJ and TRD reference fits at either length. This design removes the need to phase variants across multiple partial reads, thereby reducing chimeric allele calls caused by incorrect assignment of variants between haplotypes.

gAIRR-call version 2 was redesigned accordingly to call alleles from single Read 1 molecules. Read 2, at 100 bp too short to span a core, was used only to identify duplicate read pairs (fastp [27] version 1.3.3 with --dedup; a median of 1.8% of pairs removed per individual, range 0.06–4.95%) and was not aligned. Read 1 was quality-trimmed with fastp using a right-side sliding window of 20 bp and a mean quality threshold of 20, under two locus-aware profiles: for V loci, trimmed reads shorter than 300 bp were discarded; for D and J loci, reads were cropped to 250 bp, because adjacent TRBJ2 cores lie only 71 bp apart and longer reads extended into the neighboring gene, and trimmed reads shorter than 100 bp were discarded. Trimmed reads were aligned with minimap2 [25] to the per-gene references used for the assemblies (155 TRV, 84 TRJ and 5 TRD), with the same parameters as the assembly step. Each read was assigned to the gene against which its whole-alignment edit distance was smallest, after discounting terminal gaps introduced by the aligner’s end bonus; where alleles of two genes share a core and differ only in the flank (TRGJ1/TRGJ2 and TRBV24-1/TRBV24/OR9-2), the distinguishing flank positions had to be covered by the read and matched. A read was counted for its gene only if its sequence contained both 15-bp inner flanks of the gene, with at most two mismatches on each side (Figure 1c); the gene reference’s own flanks were tried first, and flank variants observed in the Release 1 HPRC assemblies [6] were accepted as alternative anchors (this flank table carries flank sequences only, not allele cores, and 46 of its 47 individuals are among those evaluated below). The anchor length was taken from the 15-bp seed that minimap2 uses for the V references (-k 15) and was checked on the 464 Release 2 haplotypes, each allele copy treated as an error-free read: from 12 bp upward every core was cut correctly, and longer anchors corrected no further cores while lowering the share of aligned V reads that carry the core and both anchors from 42% at 15 bp to 22% at 50 bp. The allele core was then taken from the read’s own sequence between the two anchors, and reads were grouped by that exact sequence. The support of a group was the number of molecules carrying its sequence end to end without a mismatch, each deduplicated read being treated as one molecule.

In a second pass, every first-pass group absent from the allele database and supported by at least five molecules became a provisional allele, and the reads were grouped again. A read that differed from exactly one candidate only at bases below Q20 was credited to that candidate. All candidates supported by at least five molecules first collected such reads, and a strand condition was then evaluated once for every candidate on these counts: a candidate was rejected if it was observed on a single strand with fewer than 30 molecules or, with 30 or more, if a two-sided Fisher exact test against the strand split of the largest candidate of the gene gave p < 0.05. The rejected candidates were removed, the reads were counted again, and calls were made from this final count. The strand condition targets strand-specific errors of the sequencing platform: in the 184 development samples, 99.3% of the 12,863 candidate groups that matched no assembly allele were strictly single-stranded, against 3 of the 52,325 groups that matched one; because the strand split of genuine alleles is fixed for each gene by the capture geometry, the test was made against the largest candidate of the same gene rather than against an even split. Evaluating it after masked reads have been collected lets a genuine allele whose reads from one strand all carry a low-quality error recover that strand. Per gene, groups supported by at least ten molecules were candidates, and the largest was called. Each further candidate was called if its support was at least one third of the largest; because two alleles of a gene are expected in equal numbers, this bound follows from copy number, allowing threefold imbalance, and was not tuned on accuracy. TRBV6-2 and TRBV6-3 have identical reference cores, so their reads form a single pool: the first allele of the pool draws reads from both genes, whereas a second allele carried by one of them draws reads from that gene alone, and the bound was therefore halved, to one sixth. The pool was identified from the reference sequences rather than from allele names. No limit was placed on the number of alleles per gene, so that departures from two alleles would appear as discordant calls rather than be suppressed. Genes with fewer than ten admitted reads, or with no candidate, were reported as uncalled, and these rows were kept in the output.

A called sequence identical to a database allele received its name; otherwise it was reported as its closest allele with the suffix /novel (identity ≥ 0.9), and calls from the TRBV6-2/TRBV6-3 pool were reported under a composite label. Two reporting rules resolved one-base ambiguities at the core boundary, where a flank base can be read as part of the core: a group equal to a catalogued allele plus one adjacent base of the reference flank was reported as that allele, and a group that ended or began with the reference’s boundary flank base and that, without it, equaled a catalogued allele shortened by one base was reported without that base (in the final run, the first rule never acted and the second acted 462 times in eight genes, all of whose calls were concordant). Before the per-gene conditions were applied, an edge filter discarded a novel group of the same length as its closest allele when all of its mismatches lay within 12 bp of a core end at which its representative flank was shorter than 15 bp (once in the final run, on a group arising at a D–J junction in HG002 TRBJ2-3; the gene call was concordant). For each sample and locus, gAIRR-call also reports the number of genes whose first allele has fewer than 30 molecules, below which a second allele at one third of the first would fall under the ten-molecule floor; this count was reported but not used to filter calls. All thresholds were set from first principles, from the reference and assembly sequences, or from measurements on the 184 development samples rather than by optimizing accuracy, and the gene-specific information used by the caller (distinguishing flank positions, reference bases extending beyond the canonical allele and composite labels) was derived from the reference material by rules applied to every gene.

As gAIRR-annotate does for assemblies, gAIRR-call writes the core sequences of novel alleles to a FASTA file and exports every call with its flanking sequence; because this sequence comes from reads aligned to the gene references, it extends at most 50 bp on each side, rather than the 200 bp exported from assemblies. The exported flanks are the most frequent complete pair of flanks among the reads supporting the call, masked reads included; where the two haplotypes share a core but differ in flank, the flank of the haplotype with more reads is reported, which does not affect the allele call.

### Benchmarking of targeted genotyping

Targeted genotyping was evaluated against gAIRR-annotate genotypes from the same individuals’ Release 2 assemblies, which served as ground truth (Figure 1b). gAIRR-seq version 2 libraries from 216 individuals, sequenced in four AVITI runs, were genotyped with gAIRR-call version 2: 184 development samples, on which every rule and threshold of gAIRR-call was derived; 30 held-out samples from the last sequencing run; and HG002 and HG005. The 32 samples outside the development set were genotyped once with the final program and no rule was tuned on them, although results for the 30 held-out samples had been inspected during development. HG01243 was represented by its second sequencing run, because the first was of low quality (67% of Read 1 bases at Q30 or above); the first run was analyzed separately as a test of the quality floor. Of the 216 individuals, 83 had also been sequenced with gAIRR-seq version 1 and genotyped with gAIRR-call version 1 [22]: samples already held from earlier work were sequenced with both versions, whereas newly acquired samples were sequenced mainly with version 2. Four of the 83 were excluded from the paired comparison because their two runs appear not to derive from the same library, leaving 79 individuals, all from the development set. In this paired set, the two versions were compared against the same ground truth. Because gAIRR-call version 2 relies on single reads carrying the complete core and both anchors, which 300-bp reads seldom provide for V genes [6], the two versions were compared as complete workflows, sequencing and calling together, rather than component by component.

Genotypes were compared for each individual and gene, allele by allele. The truth for a gene was the set of alleles found by gAIRR-annotate on the individual’s two haplotype assemblies, an allele carried on both haplotypes being counted once, and calls were compared with it at the sequence level, so that a called novel allele matched only if its core sequence was identical to that of the assembly allele; flanking sequence was not compared. gAIRR-call was run against material_original, which contains none of the novel alleles, so every novel allele had to be detected from the reads; novel alleles are reported under the names assigned in the material_updated rerun of gAIRR-annotate. An assembly allele reported by gAIRR-call was counted as a true positive (TP), an assembly allele not reported (including those of genes for which gAIRR-call abstained) as a false negative (FN), and a reported allele absent from the assemblies as a false positive (FP). Sensitivity was the fraction of assembly alleles that were reported, TP/(TP + FN), and precision the fraction of reported alleles that were present on the assemblies, TP/(TP + FP), each with a Wilson score 95% confidence interval. Specificity, TN/(TN + FP), took as true negatives (TN) the alleles of the gene in material_original that were neither reported nor found on the assemblies; novel alleles, being absent from the database, could only be true positives, false negatives or false positives. Alleles were counted over 243 genes per individual, with TRBV6-2 and TRBV6-3 treated as one gene because their reads cannot be separated, so that their shared core sequence was counted once. Genes with an error were classified as under-calls (an assembly allele not reported), over-calls (an additional allele reported) or misassignments (both).

Results are reported first without exclusions and then with TRDD3 set aside, the only gene excluded by name: the probe set does not capture TRDD3, and gAIRR-call abstained for it in every individual. Finally, two chromosomal segments were set aside in which independent evidence showed that one haplotype was missing from all or most cells of the cell line: the TRB 3′ segment of NA20282 (TRBV12-2 to TRBVC; a mosaic deletion) and the TRB D–J segment of HG03669 (a somatic V(D)J rearrangement). Every gene in these segments was removed together with all of its alleles, whether called correctly or not, so that the exclusion was defined by the evidence rather than by the result. Errors in genes in which raw-read pileups supported the call rather than the assembly were counted as errors. Novel allele recall was the fraction of (individual, novel allele) pairs on the assemblies that gAIRR-call reported, and a novel allele was considered present in the reads when at least ten molecules from a carrier carried its complete core without a mismatch.

### Cross-release comparison

Release 1 allele sequences and carrier assignments were obtained from the supplementary data of Yang et al. [6]. The quality-filtered Release 1 catalogue of 333 alleles was compared with the Release 2 catalogue by exact canonical-sequence matching, with containment alone not counted as rediscovery. Release 1 alleles not rediscovered were compared with the IMGT/GENE-DB reference set and classified as subsequently incorporated into IMGT when an exact match was present. For the remaining alleles, carrier information was used to determine whether any Release 1 carrier was represented in the Release 2 cohort.

### Allele frequency and population analyses

Cohort-wide allele frequency was calculated as the number of haplotypes carrying an allele divided by 464. Within each superpopulation, allele frequency was calculated as the number of haplotypes carrying the allele divided by twice the number of individuals. An allele was defined as recurrent when observed in at least two individuals and as superpopulation-private when all observed carriers belonged to a single superpopulation.

To account for unequal superpopulation sizes, each group was randomly downsampled without replacement to 31 individuals, the size of the smallest group. Superpopulation-private status was recalculated over 5,000 replicates, and the proportion of private alleles assigned to each superpopulation was summarized as mean ± SD. Allele richness was additionally rarefied to 31 individuals using the analytic hypergeometric expectation.

Analyses were performed in Python 3 using NumPy and pandas, and figures were generated with Matplotlib.

### Data availability

The sequences of the 531 novel TR alleles identified in this study are publicly accessible in FASTA format at https://zenodo.org/records/24212507, together with the sequences of all TRV and TRJ alleles observed in the 464 haplotypes and the haplotypes in which each allele was found (Supplementary Table S1). Additionally, the flanking sequence database, which includes both novel and reference alleles grouped by their 50-bp and 200-bp flanking sequences with the haplotypes carrying each group, is available at https://zenodo.org/records/24212738 (Supplementary Table S2). The gAIRR-seq sequencing reads generated in this study are not publicly available. HPRC Release 2 assemblies are publicly available through the Human Pangenome Reference Consortium [21].

## Results

### gAIRR Suite version 2 anchors allele discovery and genotyping on germline flanks

gAIRR Suite version 2 builds both of its routes on the same per-gene references, each consisting of the core of a canonical allele with 50 bp of germline flank on either side (Figure 1a). gAIRR-annotate uses the flanks to locate the core of each gene in a haplotype assembly and then compares the core with the IMGT/GENE-DB alleles of that gene, reporting a call only when the assembly supports the core together with both flanks; flanking sequence thus anchors the core without entering the allele definition. Hits whose flank diverges from the germline flank by more than 0.20 on either side, as at loci where somatic V(D)J recombination has replaced the flank with junctional sequence, are discarded before allele assignment. gAIRR-call applies the same anchoring to targeted-sequencing reads. In gAIRR-seq version 2, a 500-bp Read 1 spans a complete V core with its flanks, so that gAIRR-call version 2 calls alleles from single molecules rather than combining evidence across partial reads (Figure 1b): a read is counted only when it carries both 15-bp inner flanks, with at most two mismatches each, and the allele core is taken from the read’s own sequence between them (Figure 1c). At 300 bp, a read can meet this requirement for only 10 of the 155 TRV gene references, whereas at 500 bp it can for all of them. gAIRR-call was benchmarked against gAIRR-annotate genotypes from the same individuals’ assemblies, in the 79 individuals sequenced with both versions and in all 216 individuals sequenced with version 2 (Figure 1b).

### HPRC Release 2 identifies 531 novel TR alleles

In the 464 haplotype-resolved HPRC Release 2 assemblies [21], gAIRR-annotate version 2 found 409 of the 519 IMGT reference alleles and 536 sequences absent from IMGT/GENE-DB [18]. Five of these candidates, each carried by a single haplotype, were not observed in the targeted-sequencing reads of their carrier and were excluded (see below), leaving a catalogue of 531 novel alleles (486 TRV and 45 TRJ; Table 1; Supplementary Table S1). None of the 531 exactly matched any of the 519 reference alleles.

**Table 1.** Novel TR alleles relative to the published Release 1 catalogue and IMGT/GENE-DB.

| **Allele counts** | **Published novel<br>(2026-01)** | **IMGT<br>(2026-03-20)** |  |  |
|---|---|---|---|---|
|  |  | Total | Published novel | Strictly |
| V | 305 | 412 | 93 | 319 |
| D | 0 | 6 | 0 | 6 |
| J | 30 | 101 | 5 | 96 |
| Total | 335 | 519 | 98 | 421 |
| **Allele counts** | **Published novel<br>(2026-01)** | **V2 novel** |  |  |
|  |  | Total | Published | Strictly novel |
| V | 305 | **486** | 209 | 277 |
| D | 0 | **0** | 0 | 0 |
| J | 30 | **45** | 24 | 21 |
| Total | 335 | **531** | 233 | 298 |

Published novel (2026-01), the 335 novel alleles reported from the 47 HPRC Release 1 individuals [6]. Upper table: the IMGT/GENE-DB reference set used in this study (accessed 20 March 2026; 519 alleles), of which 98 are published Release 1 novel alleles since incorporated into IMGT/GENE-DB (Published novel) and 421 are not (Strictly). Lower table: the 531 novel alleles of the Release 2 catalogue identified with gAIRR Suite version 2 (V2 novel), of which 233 are identical to published Release 1 novel alleles (Published) and 298 had not been reported (Strictly novel); five further candidates, none previously reported, were excluded from the catalogue because they were not observed in the targeted-sequencing reads of their carriers. No novel D allele was found.

The novel alleles were distributed across all four TR loci, with 231 in TRB, 230 in TRA, 63 in TRG, and 7 in TRD. They spanned 180 genes (146 V and 34 J); TRAVA carried the most novel alleles (11), followed by TRAV4 and TRGV4 (8 each), and the median was 3 per gene (Figure 2a). The reference alleles in their names comprised 231 IMGT reference alleles, with up to eight novel alleles sharing a single reference allele (TRAV4*01 and TRGV4*02; Figure 2b); for seven novel alleles, two reference alleles were equally close and the first-listed one was used. Novel alleles were present on every haplotype, with a median of 8 per haplotype (range, 1–25) and 14 per individual (range, 5–40).

**Figure 2.**
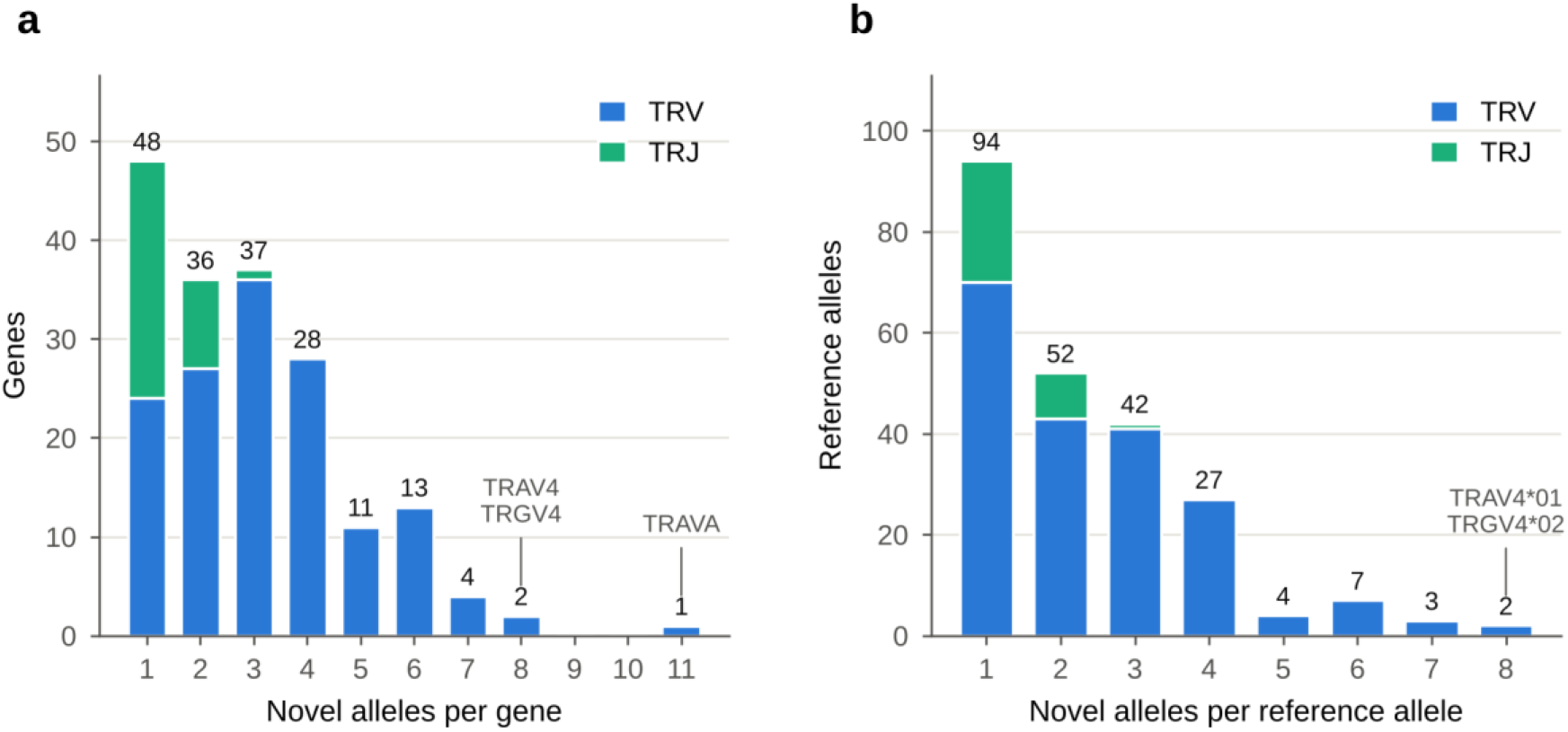
Distribution of the 531 novel TR alleles across genes and reference alleles. (a) Number of genes carrying a given number of novel alleles, for the 180 genes with at least one novel allele (146 TRV and 34 TRJ genes). (b) Number of IMGT reference alleles associated with a given number of novel alleles, counted by the reference allele in each novel allele’s name (231 reference alleles). Previously reported alleles keep their published names, so for 33 of them the named reference allele is not the closest allele in the IMGT/GENE-DB release used here; for seven alleles, two reference alleles were equally close and the first-listed one was used. Numbers above the bars are counts of genes (a) or reference alleles (b); the genes and reference alleles with the most novel alleles are named.

### Novel alleles occur in heterogeneous flanking sequence contexts

All 531 novel allele cores were recovered with both 50-bp and 200-bp germline flanking sequences. These yielded 546 distinct core-plus-50-bp-flank sequence contexts and 588 core-plus-200-bp-flank contexts. Fifteen alleles had more than one 50-bp flanking context, increasing to 43 alleles at 200 bp, showing that identical allele cores can occur within different local genomic sequence backgrounds (Supplementary Table S2).

Seven allele copies carried a flank altered by somatic V(D)J recombination in the cell line. In HG02717 haplotype 2, TRDD2 was joined to TRDD3 through an 11-nt junctional sequence (a Dδ2–Dδ3 joint), and in HG01150 haplotype 2, TRBJ2-1 was joined to TRBD1 (a Dβ1–Jβ2.1 joint); these two contexts were excluded from both the 50-bp and the 200-bp tables. Five further TRBJ2 copies lay 119–150 bp downstream of a D–J joint at the neighboring upstream J gene (TRBJ2-3 in HG002 haplotype 1 and NA19159 haplotype 2, TRBJ2-4 in HG01978 haplotype 1, TRBJ2-5 in HG03669 haplotype 2 and NA18948 haplotype 1); their 50-bp contexts are germline and were kept, and only their 200-bp contexts were excluded. Four of these joints (HG002 TRBJ2-3, HG01978 TRBJ2-4, HG03669 TRBJ2-5 and NA18948 TRBJ2-5) coincide with junction-derived J calls of gAIRR Suite version 1 (three of them reported as novel alleles), which version 2 removes by its flank-divergence filter. Two copies were left out of the 200-bp table only, because the assembly did not provide a complete 200-bp flank: an assembly gap of 81 N in the 5′ flank of a second TRAV10*01 copy in HG01243 haplotype 2, and a 1-bp deletion at the boundary between TRBV26*01 and its 3′ flank in NA18948 haplotype 2.

### Cross-release comparison accounts for all Release 1 alleles

Of the revised set of 333 Release 1 alleles [6], 233 were recovered in Release 2 by exact sequence matching. Among the remaining 100, 98 exactly matched entries now present in IMGT/GENE-DB and were therefore no longer classified as novel under the Release 2 reference baseline. The final two, TRBV5-1*01_NCWC and TRBV7-9*03_HU76, had been observed only in HG03492, the single Release 1 individual absent from Release 2, and therefore could not be reassessed. Thus, all 333 Release 1 alleles were accounted for by cross-release recovery, subsequent inclusion in IMGT/GENE-DB, or cohort composition (Table 1). The 233 recovered alleles are reported under their Release 1 names; for 33 of them the closest reference allele in the current IMGT release is one of the 98 Release 1 alleles since incorporated (for example, TRBV26*01_PF3L is now closest to TRBV26*03), and the published name was kept.

### Targeted genotyping recovers assembly alleles with high sensitivity and precision

Alleles called by gAIRR-call version 2 from gAIRR-seq version 2 reads closely matched those found by gAIRR-annotate in the corresponding assemblies (Figure 3). In the 184 development samples, on which the rules and thresholds of gAIRR-call were set, the assemblies carried 52,715 alleles in the 243 scored genes; gAIRR-call reported 52,517 of them and 5 alleles absent from the assemblies (sensitivity 99.624%, precision 99.990%). TRDD3, which the probe set does not capture, accounted for 184 of the 198 missed alleles, one in every individual; with it set aside, 14 of 52,531 alleles were missed (sensitivity 99.973%). Two chromosomal segments in which the cell line had lost one haplotype were then set aside in full: the TRB 3′ segment of NA20282, deleted in about two thirds of its cells, and the TRB D–J segment of HG03669, removed by a somatic rearrangement. This removed 58 genes carrying 75 alleles, called correctly or not, and left 6 missed and 3 extra alleles among 52,456 (sensitivity 99.989%, 95% CI 99.975–99.995%; precision 99.994%, 99.983–99.998%). The 32 samples outside the development set, on which no rule or threshold was set, gave similar results: 3 of their 9,172 alleles were missed and none was over-called (sensitivity 99.967%, 99.904–99.989%; precision 100%). Across all 216 individuals, gAIRR-call therefore missed 9 and over-called 3 of 61,628 alleles (sensitivity 99.985%, 99.972–99.992%; precision 99.995%, 99.986–99.998%): 7 missed and 3 extra among 41,642 TRV alleles, 2 missed among 19,048 TRJ alleles and no error among 938 TRD alleles; 204 of the 216 individuals had no error. Without any exclusion, the 216 individuals had 233 missed and 5 extra alleles among 61,919 (sensitivity 99.624%, precision 99.992%), 216 of the missed alleles being TRDD3. With both exclusions applied, specificity across the 216 individuals was 99.994% (95% CI 99.983–99.998%), and in gene terms the 12 errors were 12 discordant calls among 52,214 (99.977%).

**Figure 3.**
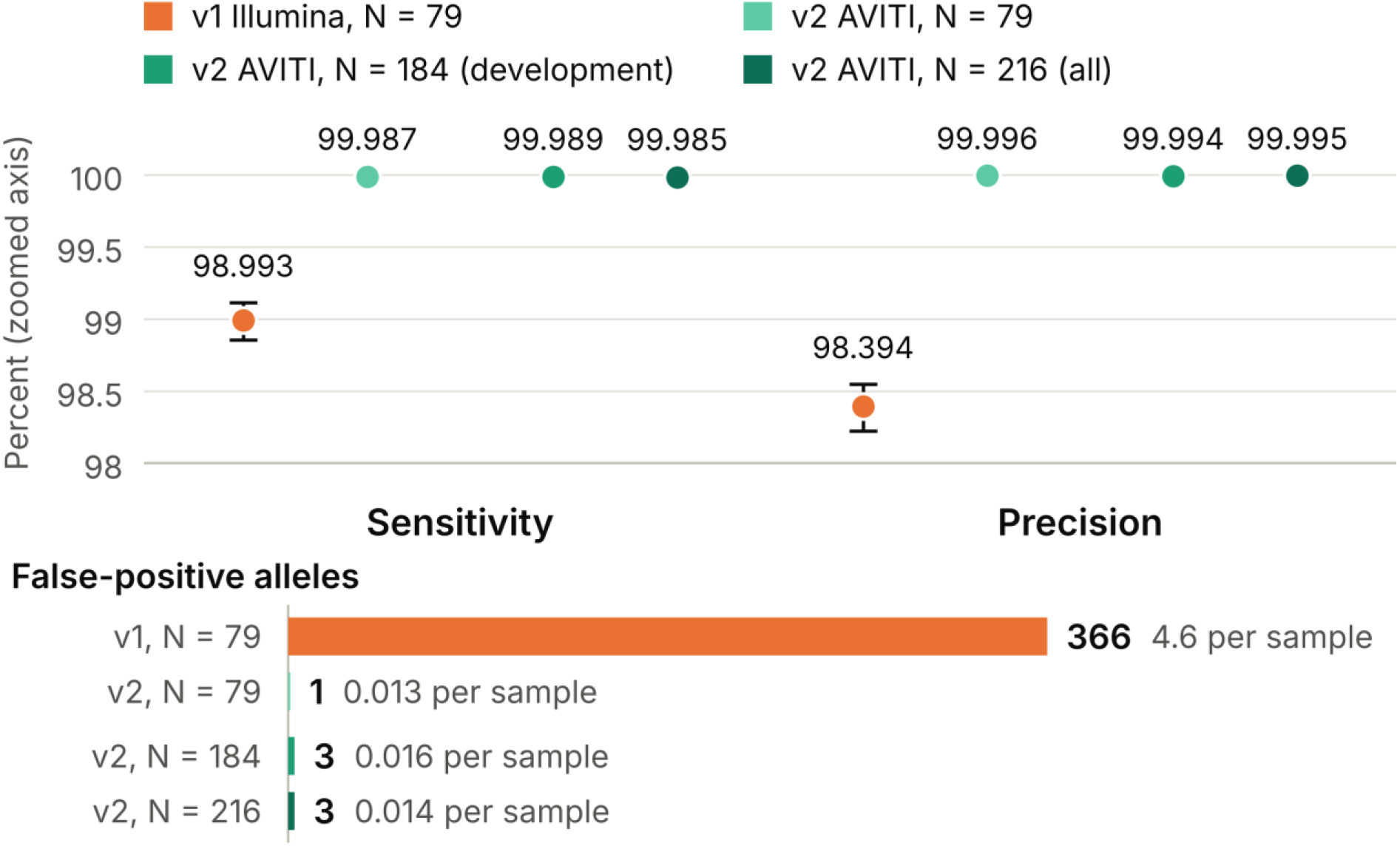
Accuracy of targeted genotyping against assembly-based genotypes. Upper panel, allele-level sensitivity and precision (zoomed axis) of gAIRR-seq and gAIRR-call version 1 (Illumina; N = 79) and version 2 (AVITI) in the same 79 individuals, in the 184 development samples and in all 216 individuals; error bars, Wilson 95% confidence intervals, which for version 2 are narrower than the symbols. Lower panel, number of false-positive alleles in each group and per sample. The ground truth is the gAIRR-annotate genotype from each individual’s Release 2 assemblies. TRDD3 is set aside in all groups and, for the 184 and 216 samples, also the two cell-line segments of NA20282 and HG03669; the 79 paired individuals are all development samples.

The 12 remaining errors, 9 missed and 3 extra alleles, were each in a different individual, and no gene call contained both a missed and an extra allele. Six were in genes in which raw-read pileups supported the call rather than the assembly, through heterozygosity collapsed in the assembly (HG01167 TRAV35, NA18570 TRBVB), junction records without read support (HG01358 TRBV7-2, HG01361 TRBJ2-7) or insertions absent from the reads (NA18944 TRBV25-1, NA18959 TRBV11-1); they were nevertheless counted as errors. One was in a gene with three copies of one allele on a haplotype (HG02886 TRGV5), so that its other allele, on one of four copies, fell below the one-third threshold (0.287 of the first allele); four were second alleles observed at 12–32% of the first, also below the threshold (HG01940 TRBVA, HG03470 TRGV10, HG00733 TRBV7-2 and HG02258 TRGJ1); and one was an over-call from a homopolymer slippage group that passed the ratio threshold (HG00673 TRAVA). Four of the nine missed alleles were novel alleles that were not observed in the targeted-sequencing reads of their carrier (TRBV7-2*01_QT7J in HG01358, TRBV25-1*01_I7A2 in NA18944, TRBV11-1*01_FZ3O in NA18959 and TRBJ2-7*01_4MN6 in HG01361); they are four of the five candidates excluded from the novel allele catalogue for this reason (the fifth, TRBJ2-7*01_HHZ6, lies in the excluded HG03669 segment), but they were kept in the truth set and counted as errors. The strand condition carried most of the filtering: of the 27,005 candidate groups ranked second or lower within a gene, 15,201 were rejected as strictly single-stranded with fewer than 30 molecules, 1,885 by the Fisher test, 358 by the one-third ratio and one by the edge filter, and 9,560 were reported as additional alleles; only one gene in one individual received a third allele.

In the 79 individuals sequenced with both versions, the assemblies carried 22,651 alleles once TRDD3 was set aside; neither excluded segment occurs in these individuals. gAIRR-seq and gAIRR-call version 1 missed 228 of these alleles and reported 366 alleles absent from the assemblies (sensitivity 98.993%, 95% CI 98.855–99.115%; precision 98.394%, 98.222–98.549%), whereas version 2 missed 3 and reported 1 (sensitivity 99.987%, 99.961–99.995%; precision 99.996%, 99.975–99.999%). The errors of version 1 lay mostly in TRV (222 missed and 353 extra among 15,333 TRV alleles, against 6 and 13 among 6,974 TRJ alleles and none among 344 TRD alleles), and in 184 gene calls version 1 both missed an assembly allele and reported a different one, which version 2 never did. Version 2 had fewer false-positive alleles than version 1 in every one of the 79 individuals, and 75 of them had no error with version 2, against none with version 1 (Figure 3).

All novel alleles of the catalogue were confirmed in the targeted-sequencing reads. Across the 216 genotyped individuals, the assemblies carried 3,497 (individual, novel allele) pairs, and gAIRR-call version 2, whose database contained none of the novel alleles, reported 3,488 of them (99.74%). Each of the 3,492 pairs involving the 531 catalogue alleles was carried by at least ten error-free molecules in the individual’s own reads; the remaining five pairs belonged to the five candidates excluded from the catalogue, none of which was observed in the reads of its carrier. Genotyping the same reads against an expanded database that contained the 536 novel candidates changed no gene call.

### Release 2 adds predominantly rare alleles

Release 2 added 298 alleles not reported in Release 1, comprising 277 TRV and 21 TRJ alleles. These additions were concentrated at low cohort frequencies. Of the 41 novel alleles with allele frequency ≥5%, 40 had been reported in Release 1; the remaining one, TRAV19*01_ON7F (71 of 464 haplotypes, 15.3%), is the allele reported in Release 1 as TRAV19*01_MTHX with four additional bases at its 5′ end. Only 11 of the 119 alleles with frequency ≥1% were absent from the Release 1 catalogue, ON7F among them. Apart from ON7F, the most frequent Release 2-specific allele, TRBV10-2*01_V32K, occurred on 11 of 464 haplotypes (2.4%). Moreover, 213 of the 298 Release 2-specific alleles (71.5%) were observed in only one individual (Figure 4).

**Figure 4.**
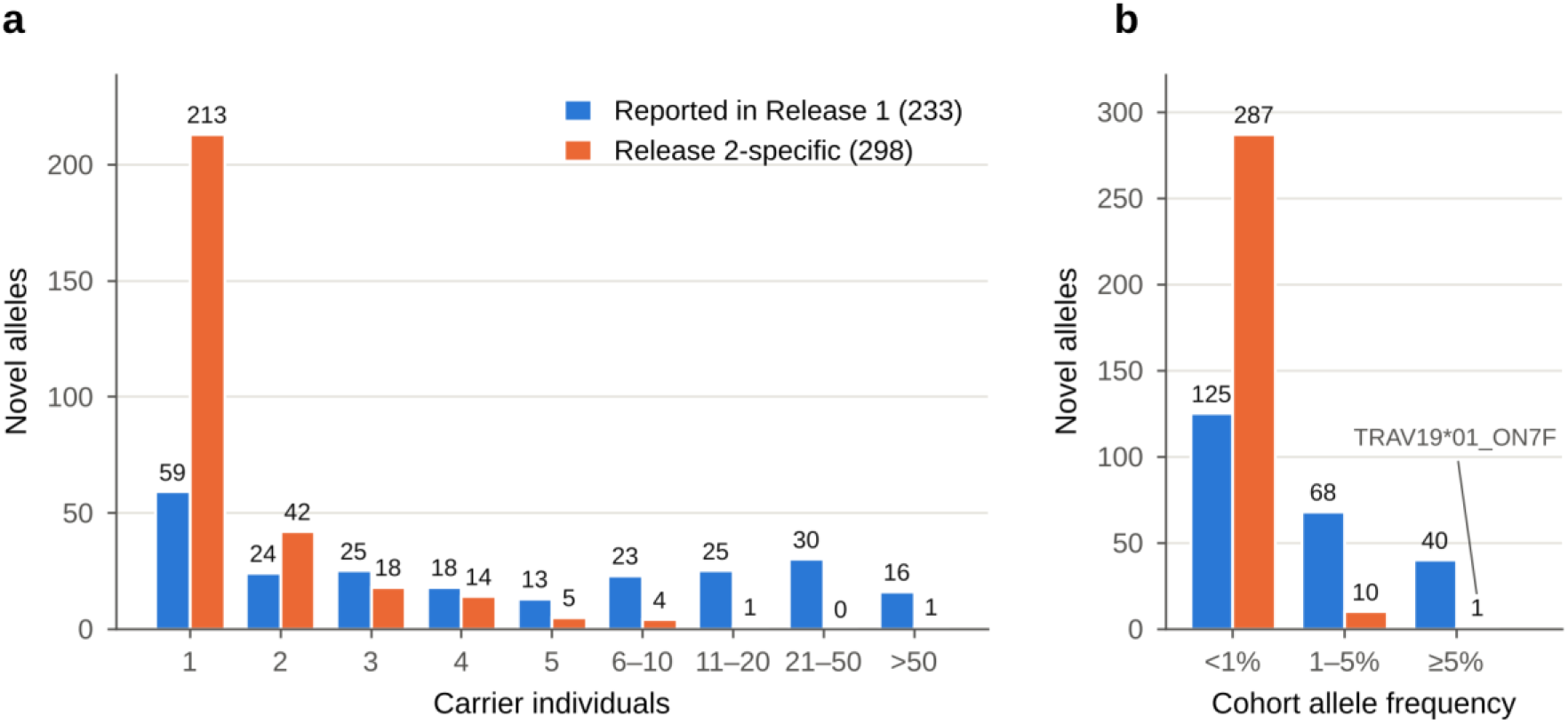
Release 2 adds predominantly rare alleles. (a) Number of carrier individuals for the 298 Release 2-specific novel alleles and the 233 novel alleles previously reported in Release 1. (b) Cohort allele frequency of the same alleles, calculated as the number of haplotypes carrying the allele divided by 464. The only Release 2-specific allele with a frequency of at least 5% is TRAV19*01_ON7F, which was reported in Release 1 with four additional 5′ bases as TRAV19*01_MTHX.

Some Release 2-specific alleles nevertheless reached higher frequencies within individual superpopulations. Besides ON7F, six had allele frequencies ≥5% in at least one superpopulation: TRBV10-2*01_V32K (9.8% in EAS), TRBV21/OR9-2*01_LV7Y (6.9% in SAS), TRGV1*01_3CON (6.5% in EUR), and TRAJ24*02_AEI2, TRAJ26*01_Q7LW and TRAV24*01_MJZE (5.9% each in EAS).

### AFR contains the largest share of superpopulation-restricted alleles

Of the 531 novel alleles, 407 were observed in only one of the five superpopulations. AFR [24] accounted for 208 of these alleles (51.1%), although it comprised 30.2% of the cohort. This skew was greater among recurrent alleles: 76.3% of the 135 superpopulation-restricted alleles observed in at least two individuals occurred in AFR (Figure 5).

**Figure 5.**
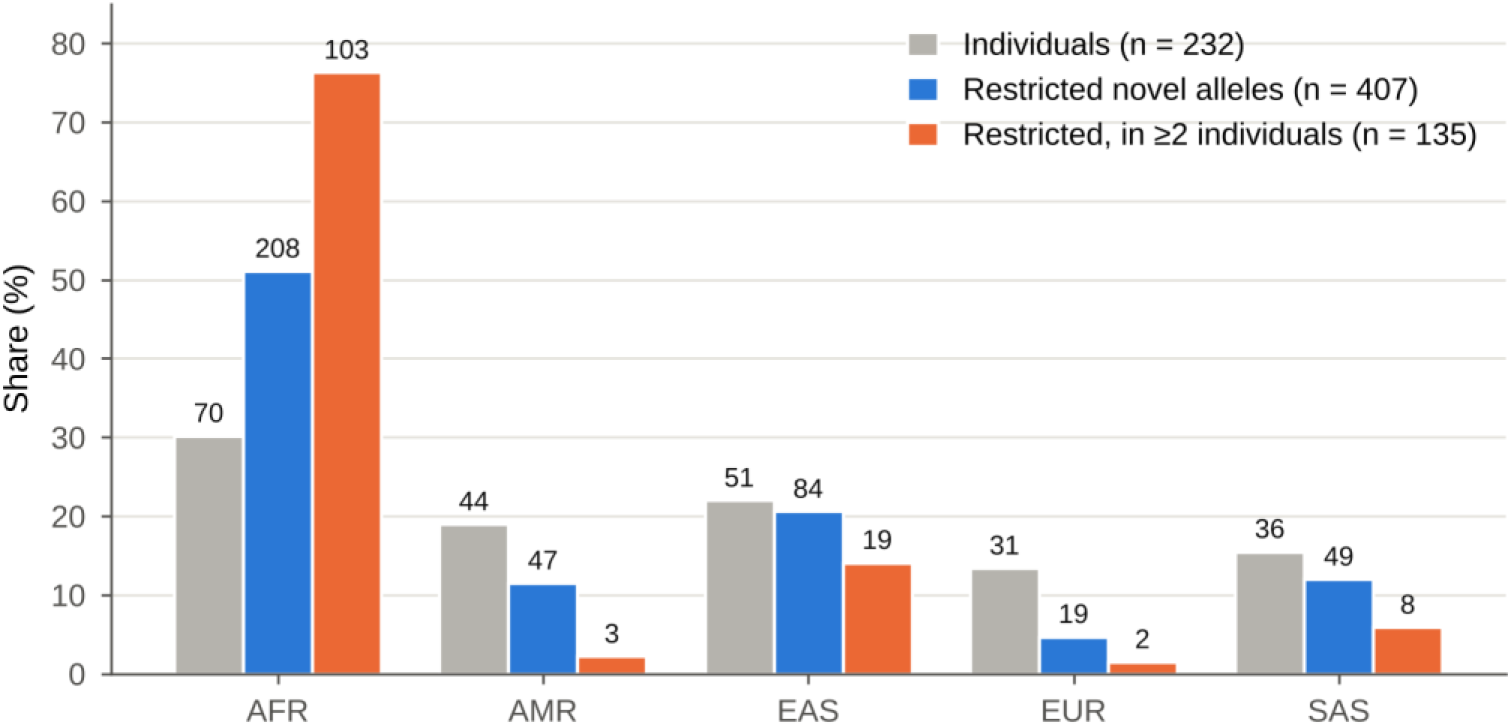
Superpopulation-restricted novel alleles. Share of each superpopulation among the 232 individuals of the cohort, among the 407 novel alleles observed in only one superpopulation, and among the 135 of these observed in at least two individuals. Numbers above the bars are counts of individuals or alleles. AFR, African; AMR, Admixed American; EAS, East Asian; EUR, European; SAS, South Asian.

The pattern persisted after equalizing sample size across superpopulations. Across 5,000 downsampling replicates of 31 individuals per group, AFR accounted for 46.0% ± 2.1% of superpopulation-restricted alleles and 67.2% ± 3.4% of recurrent superpopulation-restricted alleles, compared with an equal-share benchmark of 20%. Rarefaction to an equal sample size of 31 individuals also showed the highest novel TRV allele richness in AFR (191.0), followed by AMR (107.3), SAS (94.3), EAS (81.5), and EUR (66.0) (Figure 6).

**Figure 6.**
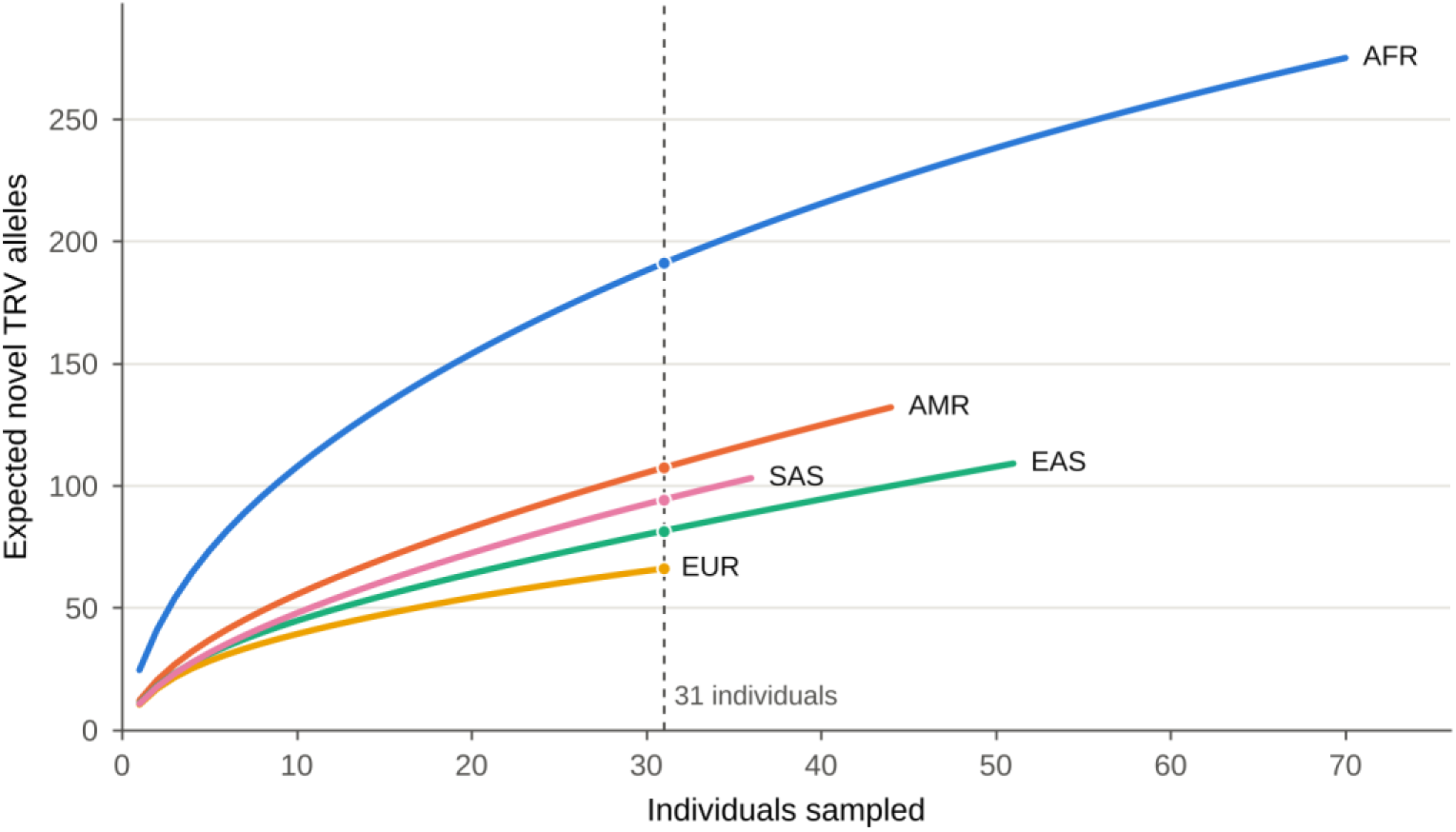
Rarefaction of novel TRV allele richness by superpopulation. Expected number of novel TRV alleles observed when n individuals are sampled from a superpopulation, calculated with the analytic hypergeometric expectation, up to the size of each superpopulation (AFR 70, AMR 44, EAS 51, EUR 31 and SAS 36 individuals). The dashed line and dots mark 31 individuals, the size of the smallest superpopulation, at which the expected numbers are 191.0 (AFR), 107.3 (AMR), 94.3 (SAS), 81.5 (EAS) and 66.0 (EUR).

## Discussion

Analysis of 232 HPRC Release 2 individuals identified 531 novel TR alleles, absent from the IMGT/GENE-DB reference set used for annotation and observed in the targeted-sequencing reads of at least one carrier, including 298 not observed in our earlier Release 1 study [6]. The expanded catalogue is supported by cross-release reproducibility, recurrence across individuals and targeted-capture genotyping. Retrospective reanalysis of the Release 1 catalogue under the flanking-support criteria introduced in gAIRR Suite version 2 withdrew two of the 335 alleles reported in that study, leaving 333 high-confidence alleles as the baseline for this comparison. Of these, 233 were rediscovered in Release 2 and 98 have since been incorporated into IMGT/GENE-DB. The remaining two could not be reassessed because their only carrier was absent from Release 2. All 333 are accounted for by rediscovery, reference adoption, or cohort composition.

Despite an approximately five-fold increase in cohort size, Release 2 yielded few additional common alleles. Among the 41 novel alleles with a frequency of at least 5%, only TRAV19*01_ON7F had not been reported in Release 1, where the same allele had been reported with four additional 5′ bases as TRAV19*01_MTHX, and even at a 1% frequency threshold only 11 of 119 alleles were new. By contrast, 213 of the 298 Release 2-specific alleles were singletons. Low frequency across the full cohort, however, does not reflect low frequency within individual populations. When frequencies were calculated separately within each superpopulation, six Release 2-specific alleles besides ON7F reached a frequency of at least 5%. For example, TRBV10-2*01_V32K reached 9.8% in East Asian individuals (10 of 102 haplotypes). Release 2 primarily added alleles that are globally uncommon yet locally frequent, highlighting the importance of diverse population representation for expanding the TR germline reference.

The population-specific enrichment was most prominent in African-ancestry individuals. Of the 407 alleles observed in only one superpopulation, 208 (51.1%) were found exclusively in African-ancestry individuals, although they comprised only 30.2% of the cohort. When the analysis was restricted to alleles observed in at least two individuals, 76.3% of these superpopulation-private alleles were found in African-ancestry individuals, indicating that the pattern was not driven primarily by singletons. Because the African ancestry group was the largest, we repeated the analysis after downsampling each superpopulation to 31 individuals. Across 5,000 downsampling replicates, African-ancestry individuals still accounted for 46.0% of superpopulation-private alleles and 67.2% of recurrent private alleles. Thus, differences in group size alone cannot explain the observed enrichment. Greater genetic diversity in African populations likely contributes to this pattern [7], while their historical underrepresentation in germline reference resources may also cause more existing variation to be classified as novel [10]. We cannot distinguish the relative contributions of these two factors, but both support broader population representation in future TR germline reference development.

These population-level patterns are consistent with the recent large-scale study by Mantena et al. [7], who used long-read data from 2,719 donors, including 227 HPRC Release 2 individuals, and identified 491 novel V-gene and 79 novel J-gene alleles. The two studies have distinct but connected scopes. Mantena et al. focused on defining common TR variation and excluded pseudogene-class segments, whereas our catalogue retains pseudogene-class alleles and singletons and provides 200 bp of genomic sequence on both sides of each allele. Their larger cohort and rarefaction analysis suggest that much of the common TR allele diversity has already been captured. Our results, by contrast, show that increasing population representation continues to uncover rare alleles that are enriched within specific populations. The two studies therefore converge on the need for population-diverse germline references while providing complementary information on common variation, rare alleles, and local genomic context.

Assembly-based and repertoire-based approaches provide different vantage points on TR germline variation. Repertoire-based inference identifies alleles from rearranged receptor sequences, providing direct evidence that an allele is rearranged and expressed. Using this approach, Corcoran et al. identified 175 novel V- and J-gene alleles in 45 deeply sequenced donors [28], and Mikelov et al. identified 81 novel alleles in 134 individuals [29]. Such inference cannot, however, recover alleles absent from the sampled repertoire or characterize pseudogene-class segments and their surrounding genomic sequence. Assembly-based analysis instead surveys the inherited locus independently of expression while preserving genomic context, which serves both biological and technical purposes in our framework. Biologically, the 200-bp flanks enable direct examination of RSS and nearby regulatory variation [6]. Technically, gAIRR Suite version 2 incorporates 50 bp of germline sequence on each side of the allele core and reports a candidate only when both flanks are supported by the assembly, and applying this stricter criterion led to the withdrawal of two Release 1 calls described above. One of these, TRBJ2-3*01_FA4L, was carried by a single haplotype whose 5′ flank diverged from the germline flank by 0.62, consistent with somatic rearrangement rather than a germline allele. These complementary approaches prioritize either direct evidence of repertoire usage or comprehensive characterization of the underlying germline locus. At larger cohort sizes, short-read methods provide an additional route for extending established germline catalogues to population-scale genotyping. Approaches such as pmTR [30] and gAIRR-wgs [31] substantially increase scalability, although highly homologous and duplicated TR genes remain difficult to resolve.

Important uncertainties remain in the calls themselves. The predominance of singletons among Release 2-specific alleles makes genuine rare variants difficult to distinguish from local assembly errors. This concern is supported by our Release 1 analysis, in which several candidates were subsequently traced to assembly artifacts. Independent targeted-capture genotyping is therefore particularly important for validating singleton alleles. gAIRR-seq and gAIRR-call version 2 provide a more stringent, assembly-independent validation framework by requiring direct support for the complete allele sequence from genomic DNA and by avoiding ambiguous evidence from highly similar paralogues. Importantly, the revised pipeline preferentially withholds calls when these criteria are not met rather than reporting weakly supported alleles. This more conservative design reduces false-positive calls and should improve confidence in experimentally validated novel alleles.

The caller’s floor of ten exact molecules and its one-third heterozygosity ratio imply that a second allele can be missed whenever the first allele of a gene is supported by fewer than 30 molecules. In the one library sequenced below the usual quality (HG01243, first run: 67% of Read 1 bases at Q30 or above and 1.45 million read pairs), the final caller missed the second allele of two V genes, both among the genes whose first allele had fewer than 30 molecules, and over-called none; re-sequencing the library (87% at Q30 or above, 2.10 million read pairs) resolved both. The number of genes below the 30-molecule mark is reported per sample and locus; for TRV it was zero in 182 of the 184 development samples.

Beyond uncertainty in individual allele calls, the population-level interpretation of the catalogue also remains provisional. The broad superpopulation labels used here do not capture finer population structure or admixture, and 232 individuals remain insufficient to determine whether the long tail of rare TR variation has been comprehensively sampled. The catalogue should therefore be viewed as a substantial expansion of the human TR germline reference, rather than an exhaustive representation of its population diversity.

Despite these limitations, a more complete reference measurably improves downstream genotyping. In gAIRR-wgs, TRV recall declined when more than 10% of novel alleles were removed from the reference, whereas TRJ genotyping was more tolerant of reference depletion [31]. Reference breadth also determines which alleles appear population-specific. Six alleles that seemed specific to the Taiwan Biobank when compared with the 47 Release 1 individuals were subsequently found in East or South Asian individuals after Release 2 expanded the comparison set [31]. As a result, larger and more diverse reference cohorts can reduce the risk of classifying an allele as population-specific simply because other populations were insufficiently sampled. As these references mature, the next step is to move from allele discovery toward functional and population-scale analysis. This will require harmonized allele definitions across long-read catalogues, independent validation of uncertain alleles, matched repertoire data to determine which alleles are rearranged and expressed, and scalable genotyping or imputation for association studies. Similar studies of immunoglobulin loci have shown that inherited germline variation can shape repertoire usage [32], providing a precedent for investigating analogous effects of TR germline variation.

Overall, expanding the HPRC cohort from 47 to 232 individuals added relatively few common TR alleles but substantially increased the detection of rare alleles, including variants enriched within specific superpopulations. Thus, the main benefit of population-scale pangenome analysis is not simply finding more alleles, but identifying variants that would be missed in smaller or less diverse cohorts. As pangenome resources expand, the focus can increasingly shift from allele discovery toward determining how inherited TR variation influences receptor usage, immune phenotypes, and disease.

## Supplementary tables

**Supplementary Table S1. Sequences of novel alleles.** TRV and TRJ alleles observed by gAIRR-annotate in 464 haplotype-resolved HPRC Release 2 assemblies (232 individuals), including the 531 novel alleles of the catalogue (486 TRV and 45 TRJ; previously published novel alleles keep their published names). allele, allele name; haplotype, the haplotypes carrying the allele (h1 and h2, haplotype 1 and 2 of the HPRC Release 2 assembly; a haplotype with two or more copies of the allele is listed once); haplotype_count, their number; seq, nucleotide sequence of the allele.

**Supplementary Table S2. Flanking sequence database.** Groups of TR alleles by their 50-bp and 200-bp flanking sequences in 464 HPRC Release 2 haplotypes (232 individuals). seq_id_cluster, allele and group number; haplotype, the haplotypes carrying the group (h1 and h2, haplotype 1 and 2 of the HPRC Release 2 assembly); haplotype_count, their number; sequence, flank + allele + flank.

## Notes

https://zenodo.org/records/24212507

https://zenodo.org/records/24212738

### Competing Interest Statement

The authors have declared no competing interest.

